# Global population genomic analysis of *Mycoplasma bovis* isolates reveals transcontinental variations and potential virulence genes

**DOI:** 10.1101/2020.08.19.257345

**Authors:** Roshan Kumar, Karen Register, Jane Christopher-Hennings, Paolo Moroni, Gloria Gioa, Nuria Garcia-Fernandez, Julia Nelson, Murray Jelinski, Inna Lysnyansky, Darrell Bayles, David Alt, Joy Scaria

## Abstract

Among more than twenty species belonging to the class Mollecutes, *Mycoplasma bovis* is the most common cause of bovine mycoplasmosis in North America and Europe. Bovine mycoplasmosis causes significant economic loss in the cattle industry. The number of *M. bovis* positive herds has recently increased in North America and Europe. Since antibiotic treatment is ineffective and no efficient vaccine is available, *M. bovis*-induced mycoplasmosis is primarily controlled by herd management measures such as the restriction of moving infected animals out of the herds and culling of infected animals or shedders. To better understand the population structure and genomic factors that may contribute to its transmission, we sequenced 147 *M. bovis* strains isolated from four different countries and hosts, primarily cattle. We performed a large-scale comparative analysis of *M. bovis* genomes by integrating 104 publicly available genomes and our dataset (251 total genomes). A whole genome-single nucleotide polymorphism (SNP)-based phylogeny revealed that *M. bovis* population structure is composed of five clades with one of the isolates clustering with the outgroup *M. agalactiae*. These isolates were found to cluster with those from Canada, Israel, Lithuania, and Switzerland, suggesting trans-continental transmission of the strains. We also validated a previous report suggesting minimum divergence in isolates of Australian origin, which grouped within a single clade along with strains from China and Israel. However, no observable pattern of host association in *M. bovis* genomes was found in this study. Our comparative genome analysis also revealed that *M. bovis* has an open pangenome with a large breadth of unexplored diversity of genes. Analysis of *vsp* gene-host association revealed a single *vsp* significantly associated with bovine isolates that may be targeted for diagnostics or vaccine development. Our study also found that *M. bovis* genome harbors a large number of IS elements, including a novel 1624 bp IS element, and ISMbov9. Collectively, the genome data and the whole genome-based population analysis in this study may help to develop control measures to reduce the incidence of *M. bovis*-induced mycoplasmosis in cattle and/or to identify candidate genes for vaccine development.

## Introduction

*Mycoplasma bovis* is a member of the class Mollicutes, representing the simplest, wall-less, self-replicating bacterium, known to cause multiple diseases including pneumonia, arthritis and confer late-term bovine abortions [1]. During the course of evolution, the genus *Mycoplasma* underwent regressive evolution and maintained the minimal set of biosynthetic genes necessary for its survival and adjusted to the parasitic lifestyle [2]. *M. bovis* can infect a wide variety of hosts such as cattle, bison, sheep and goats and shows tissue specificity within these hosts [3]. The first *M. bovis* strain was isolated in the US in 1961 from milk obtained from a dairy herd severely affected with mastitis[4], and it is speculated that the subsequent transportation of infected animals led to its spread across China, Australia, Israel and other European countries [5]. This pathogen has been reported to successfully evade the host immune system through the attachment mediated by variable surface proteins (Vsps) along with the biofilm formation [6, 7]. For these reasons, *M. bovis* is a significant threat to animal health worldwide and is responsible for substantial economic loss [8, 9]. In order to control *M. bovis*, the New Zealand government recently approved eradication of large number of cows with the hopes of eliminating *M. bovis* from the island nation, causing an economic loss of hundreds of millions of dollars [10]. Worldwide, *M. bovis* still remains a threat to the dairy industry even after 59 years since its first isolation.

Altogether, it has been well established that the *M. bovis* infection is a multifactorial process and adhesion is the key step towards infection and colonization, mainly mediated through Vsps, which have the ability to undergo substantial antigenic variation involving high-frequency phenotypic switching [11, 12]. Therefore, to understand the extended genetic repertoire of Vsps and to gain deep insight into the phylogeny and genomic attributes, we performed a comparative genomic analysis of 251 *M. bovis* strains, representing the largest genomic dataset studied so far. To this end, we integrated 104 *M. bovis* genomes from the public repositories to our dataset of 147 *M. bovis* isolated between 1994 and 2019 from four different countries and four different hosts. This study provides detailed insight into the phylogenetic relationships, virulence profiles, insertion sequences and functional attributes of the *M. bovis* isolates.

## 2. Materials and methods

### Genomes included in the comparative analysis

Altogether, we sequenced 147 *M. bovis* isolates from four different countries and four different hosts. We downloaded 104 publicly available *M. bovis* genomes from NCBI for comparative analysis with our dataset. Therefore, the total dataset used for comparative analysis is 251 genomes (Supplementary Table 1). The strains sequenced in this study were grown in pleuropneumonia-like organism (PPLO) medium supplemented with 10% horse serum (Thermo Fisher Scientific, Waltham, MA, USA) at 37°C for 48–72 h. The genomic DNA was isolated using DNeasy Blood & Tissue kit (Qiagen, Germany) according to the manufacturer instructions and quantified using Qubit Fluorometer 3.0 (Invitrogen, Carlsbad, CA). The whole genome sequencing was performed by the Illumina MiSeq sequencer using paired-end V3 chemistry. The datasets for six isolates named NADC83, NADC81, NADC72, NADC53, NADC51 and KRB1 were obtained from the National Animal Disease Center, Ames, IA. These isolates were sequenced using both PacBio and Illumina platforms. PacBio sequencing was carried out by the Yale Center for Genome Analysis (New Haven, CT). Following fragmentation and end repair of genomic DNA, BluePippen size selection was used to enrich for 10-20 bp fragments. Libraries were sequenced using a single SMRTcell per isolate on a PacBio RS II instrument using P6-C4 chemistry. Illumina sequencing (MiSeq; Illumina, San Diego, CA) was carried out at the National Animal Disease Center using 2 × 150 paired-end libraries, prepared with a Nextera XT DNA library preparation kit (Illumina), as detailed in the Reference Guide. The MiSeq 2 × 300-bp paired-end reads for 16 strains (NADC5, NADC15, NADC24, NADC28, NADC30, NADC32, NADC44, NADC45, NADC50, KRB5, KRB6, KRB7, KRB8, KRB9, KRB10 and KRB11) were provided by Dr. Murray Jelinski. Thereafter, we included eight complete *M. bovis* genomes (08M, Ningxia-1, CQ-W70, HB0801, Hubei-1, NM2012, JF4278 and PG45) from the NCBI database. We also downloaded genome datasets for 95 *M. bovis* strains from the Sequence Read Archive (SRA), based on the availability of metadata information on or before March 31^st^, 2019 (Supplementary Table 1). Therefore, a total of 251 *M. bovis* genomes were included in this analysis. The outgroup *M. agalactiae* strain 5632 was selected based on genome distance calculation using pairwise average nucleotide identity (ANI) [13].

### Genome assembly and validation

The raw reads were assembled into contigs using Unicycler [14] that builds an initial assembly graph from short reads using SPAdes 3.11.1 [15], followed by read correction using pilon [16]. The advantage of using Unicycler is that it can also assemble a combination of short and long reads, with high levels of accuracy. The assembled contigs (>500bp) were validated using QUAST [17]. All assemblies include 200 or fewer contigs and have an N50 of ≥10,000.

### Genome annotation, core and pangenome analysis

All these assembled genomes were annotated using Prokka [18]. A manually annotated reference *M. bovis* (PG45, Hubei-1, HB0801, CQ-W70, NM2012) genbank file was downloaded (https://www.ncbi.nlm.nih.gov/genome/browse#!/prokaryotes/mycoplasma%20bovis) and formatted to a prokka database file format. Open reading frames (ORFs) were predicted and annotated using prokka (-gcode 4) against the formatted database [18]. The general feature format (.gff) files from prokka were used as input for Roary pipeline (percentage identity ≥ 90) [19] to generate the core genome alignment using PRANK [20]. This was then used to predict the polymorphic sites using snp-sites package [21]. Then, the output was used for model testing using the package modeltest-ng (https://github.com/ddarriba/modeltest). The tree was constructed using Generalized Time Reversible (GTR) + G4 model in RAxML [22]. The pangenome matrix generated in Roary was parsed using R (R Core Team, 2018) considering the prevalence of genes with a distribution 5% < x < 99% for better clarity and plotted with core gene phylogeny.

Further, the orthologs were redefined using orthoMCL [23] available in GET_HOMOLOGUES software package given the following parameters: percentage identity ≥ 90% and query coverage of ≥75%, using GenBank files. The orthologs thus obtained were annotated using the RAST server using genetic code 4.

### Phylogeny reconstruction and functional analysis

In this study, we inferred phylogeny based on whole genome SNPs using kSNPv3.0 [24] with a k-mer length of 31. The SNP-matrix file was used for model testing using the package modeltest-ng (https://github.com/ddarriba/modeltest). GTR + G4 was the best scoring model predicted and this substitution model was used in RAxML [22] to construct the maximum-likelihood (ML) tree. The phylogenetic tree was visualized using GrapeTree [25].

To functionally characterize the *M. bovis* genomes, the amino acid sequences were searched against the eggNOG database [26]. The length of query protein was set to at least 50% in order to be involved in the alignment. The resulting KEGG annotations (KO identifiers) assigned to amino acid sequences of individual isolates were parsed in R (R Core Team, 2018) to generate the abundance matrix. This matrix was used in MeV to construct the gene tree using Pearson correlation and hierarchical clustering [27]. This was then visualized using interactive Tree of Life (iTOL) [28].

### Identification of host-associated genes

We applied Scoary to identify genes significantly associated with particular traits, such as host species [29]. This package uses gene presence/absence matrix from Roary and combines Fisher’s exact test, a phylogeny-aware test and an empirical label-switching permutation analysis. The genes that were most significantly associated with a trait, either positively or negatively, were sorted based on *p* adjusted values using multiple test correction (*p* <0.05) and their nucleotide sequences used to query the NCBI database to assign proper gene function [30].

### Identification of *vsp* genes and insertion sequences (IS)

To identify *vsp* genes, we used a 70 bp highly conserved nucleotide sequence found immediately 5’ of the start codon in all *vsp* genes as a reference and formatted the database. All the genomes were then searched against this database using the parameters: query coverage 85% and percentage identity 85%.

For IS elements, we extracted the Prokka-annotated IS element sequences (n=2640) present in 251 *M. bovis* isolates. Along with this, we included the ISfinder database sequences (n=5685) updated on July 25^th^, 2018. Cd-hit was then used to cluster the sequences using 50% query coverage and 95% sequence identity [31]. At the end, 4878 sequences were used to format the database. Individual genomes were searched against the formatted database using 70% query coverage and 95% sequence identity to identify novel IS elements. Finally, the comparative abundance matrix of IS elements was generated for the *M. bovis* isolates and visualized in R using the package ComplexHeatmap (R Core Team, 2018).

## Results

### Genome sequencing and general genomic attributes of geographically diverse *M. bovis* isolates

In this study, we included the sequence of 147 *M. bovis* strains isolated between 1994 and 2019 from four different countries viz., USA (n=121), Canada (n=22), Israel (n=3) and Lithuania (n=1) (Supplementary Table 1). These strains were mainly contributed by two host types namely bovine (n=75) and bison (n=70). Out of 147 *M. bovis* strains, four (NADC 51, NADC72, NADC 81 and NADC83) were assembled into single contigs. We then added eight complete and 95 reassembled draft *M. bovis* genomes from the NCBI database, for which at least partial metadata information was available (Supplementary Table 1). Altogether, a total of 251 *M. bovis* genomes were analyzed in this study, by far the largest whole genome dataset for *M. bovis* isolates. Based on origin, these isolates were distributed across seven countries: the United States (56%), Australia (30.8%), Canada (8.8%), China (2.4%), Israel (1.2%), Lithuania (0.4%) and Switzerland (0.4%) (Table 1). Bovine genomes constituted the largest dataset representing 71.2% (n= 179) of genomes, followed by bison isolates (28%; n=70) and one isolate each from mule deer and white tail deer. Most of these isolates were isolated from either the lung (36.05%, n=53) or milk (22.45%, n=33).

**Table 1:**
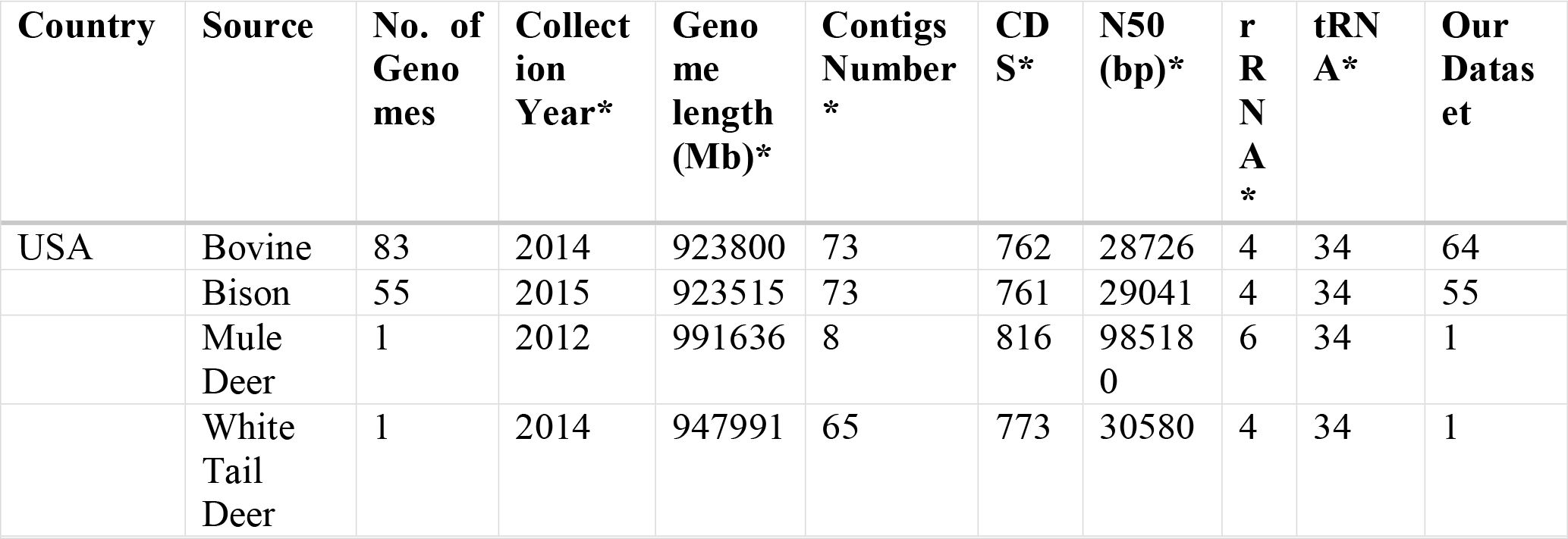
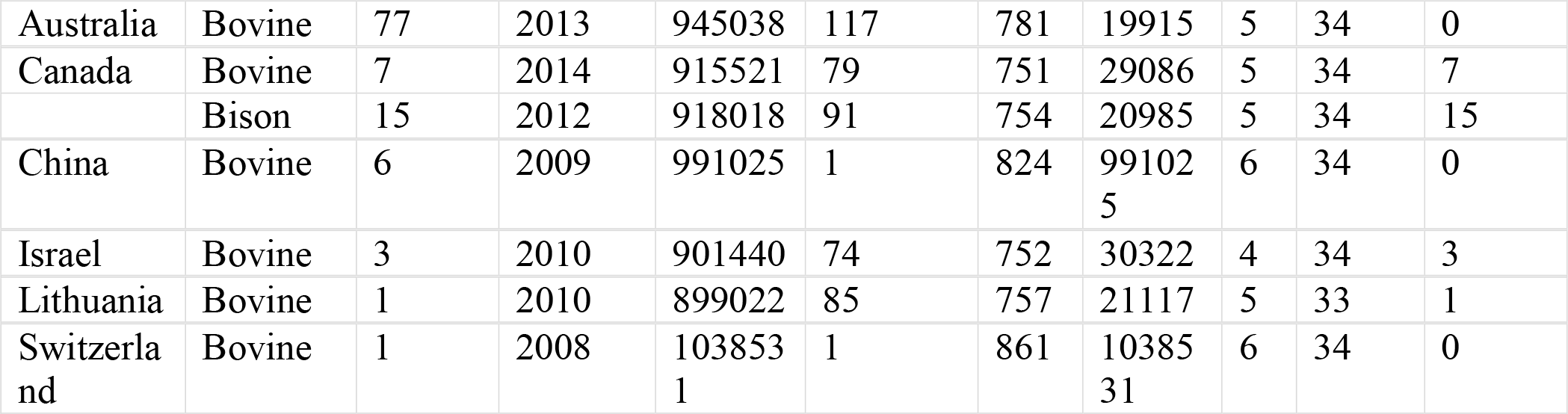
The metadata representation of 251 *M. bovis* isolates used in the study (*Median).

| Country | Source | No. of<br>Genomes | Collect<br>ion<br>Year* | Genome<br>length<br>(Mb)* | Contigs<br>Number<br>* | CD<br>S* | N50<br>(bp)* | r<br>R<br>N<br>A<br>* | tRNA* | Our<br>Dataset |
| --- | --- | --- | --- | --- | --- | --- | --- | --- | --- | --- |
| USA | Bovine | 83 | 2014 | 923800 | 73 | 762 | 28726 | 4 | 34 | 64 |
|  | Bison | 55 | 2015 | 923515 | 73 | 761 | 29041 | 4 | 34 | 55 |
|  | Mule<br>Deer | 1 | 2012 | 991636 | 8 | 816 | 98518<br>0 | 6 | 34 | 1 |
|  | White<br>Tail<br>Deer | 1 | 2014 | 947991 | 65 | 773 | 30580 | 4 | 34 | 1 |
| Australia | Bovine | 77 | 2013 | 945038 | 117 | 781 | 19915 | 5 | 34 | 0 |
| Canada | Bovine | 7 | 2014 | 915521 | 79 | 751 | 29086 | 5 | 34 | 7 |
|  | Bison | 15 | 2012 | 918018 | 91 | 754 | 20985 | 5 | 34 | 15 |
| China | Bovine | 6 | 2009 | 991025 | 1 | 824 | 991025 | 6 | 34 | 0 |
| Israel | Bovine | 3 | 2010 | 901440 | 74 | 752 | 30322 | 4 | 34 | 3 |
| Lithuania | Bovine | 1 | 2010 | 899022 | 85 | 757 | 21117 | 5 | 33 | 1 |
| Switzerland | Bovine | 1 | 2008 | 1038531 | 1 | 861 | 1038531 | 6 | 34 | 0 |

The assembly statistics revealed an average genome size of 0.93 ± 0.034 Mbp; with largest genome size in case of strain NADC53 and smallest in case of strain VDL100 (range: 0.839 to 1.071 Mbp). The average GC content was 29.49%, with a maximum and a minimum of 30.27% for Mb61 and 29.25% for NADC53, respectively. The average number of coding sequences was 770 (range: 882-694). When complete genomes of *M. bovis* were analyzed, we find that there as many as 68 ISs with a total size of approximately 0.085 Mbp/genome, suggesting a high level of genomic rearrangement within the genomes of *M. bovis* isolates (Supplementary Table 1).

### Phylogenetic structure of *M. bovis* isolates

Widely used methods to determine the evolutionary relationship among members of the genus *M. bovis* are multilocus sequence typing (MLST) and multiple-locus variable-number tandem repeat (MLVA) [32-34]. In this study, we implemented whole genome single nucleotide polymorphism (SNP) method to infer the phylogeny. To resolve this, we used *M. agalactiae* strain 5632 as an outgroup, because this species is the closest (<83.85% at ANI level) relative of *M. bovis*. Using the SNP-based phylogeny, the *M. bovis* isolates formed six different clades (Figure 1). Based on geographical location, the Australian isolates formed a separate clade (clade VI) along with all the Chinese (CQ-W70, Hubei-1, NM2012, 08M, HB0801 and Ningxia-1) and two Israeli (strain 78204 and 88172) isolates (Figure 1). In contrast to this, the USA isolates showed a high degree of genomic divergence and thus clustered in five different clades (clade I-V). Out of five clades, four (I, III, IV and V) were exclusively occupied by the USA isolates (Figure 1). This could be because of sampling bias as nearly 56% of the isolates originated from the USA. However, 15 Canadian isolates and one each from Switzerland (JF4278), Israel (87793) and Lithuania (F148) clustered with the USA isolates in clade II (Figure 1). For *M. bovis* isolates, strain CG1-1544 from clade I clustered with the outgroup *M. agalactiae* strain 5632, showing the highest degree of ancestral relationship. Clades III and V were dominated by bison isolates, but there was not a clear-cut clustering pattern based on host origin. The isolate from white tail deer (KRB8), clustered with bovine isolates in clade II. In contrast to this, a mule deer isolate (KRB1) did not fall in any major clade, but instead clustered with the bison isolates NADC98 and NADC97. There is no apparent epidemiologic link between KRB1, a 2012 isolate from Nevada, and NADC97 and NADC98, which were obtained in 2017 from different anatomic sites of a single bison in Alberta.

**Figure 1:**
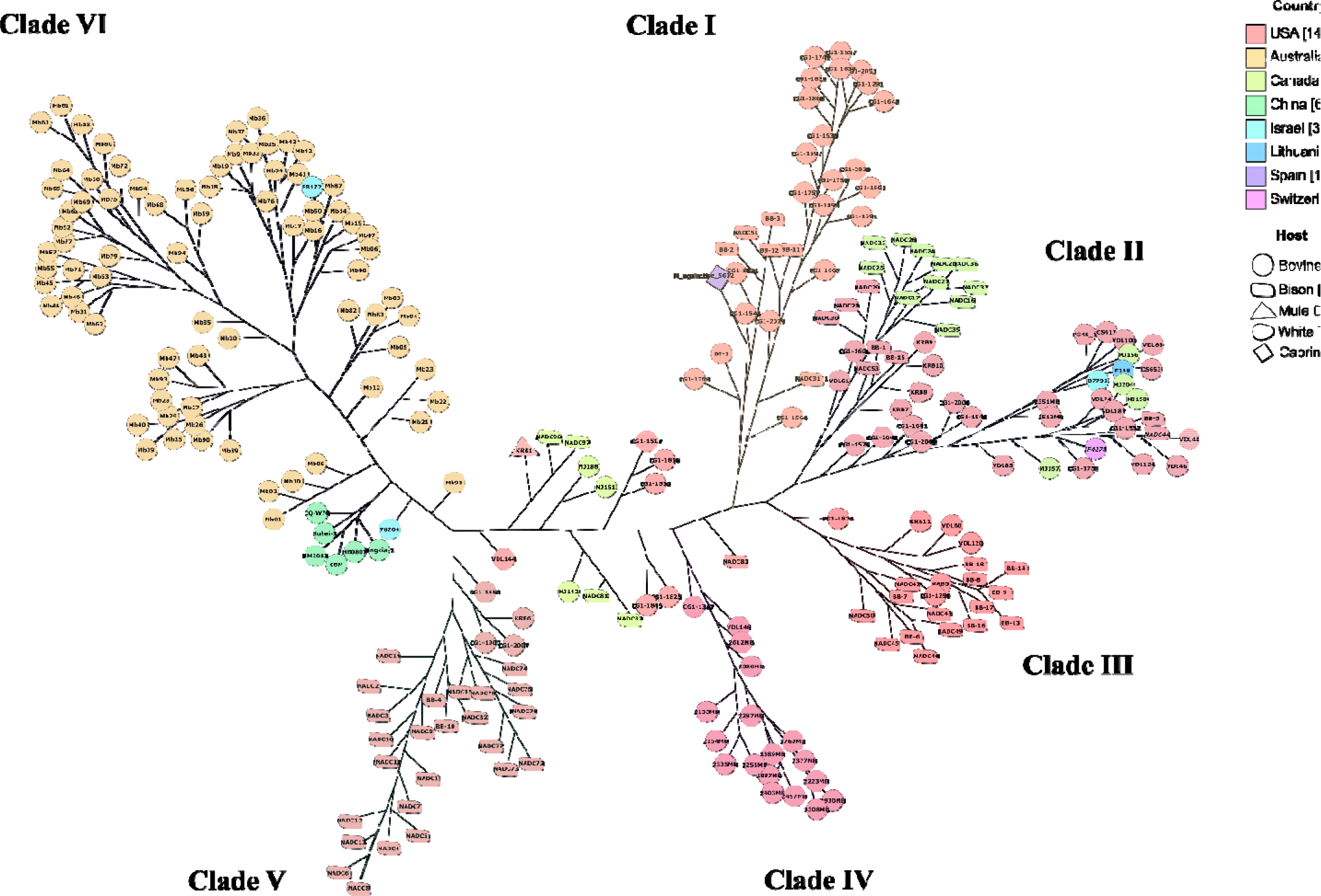
Whole-genome SNP-based phylogenetic tree of 250 *M. bovis* strains rooted to *M. agalactiae* 5632 generated by kSNP3.0. The SNP-matrix file was used for model testing using the package modeltest-ng. Thereafter, the maximum likelihood tree was constructed using Raxml-ng (GTR+G4 model). The tree was visualized using GrapeTree. The color code represents their country of isolation.

### Core and pangenome analysis

The core and pangenome analysis revealed the presence of 283 and 1186 coding genes across 250 *M. bovis* isolates, respectively. The high number of accessory genes observed in pangenome suggested that the pangenome is still not fully complete and the addition of more *M. bovis* genomes will increase the pangenome size (Figure 2). The conserved 283 coding sequences span across 40 subsystems with an average size of 0.248Mb. A majority of these genes code for ribosomal (n=45; 15.9%) and hypothetical (n=51; 18.02%) proteins. However, the core genome was also found to harbor genes associated with virulence. A 454-amino acid α-enolase (phosphopyruvate hydratase) protein was conserved in the core genome of *M. bovis*. Two lipoate-protein ligase A (*lpl*A) genes were also identified in the core genome. Interestingly, *lplA* genes were also identified as potential virulence factors [35, 36].

**Figure 2:**
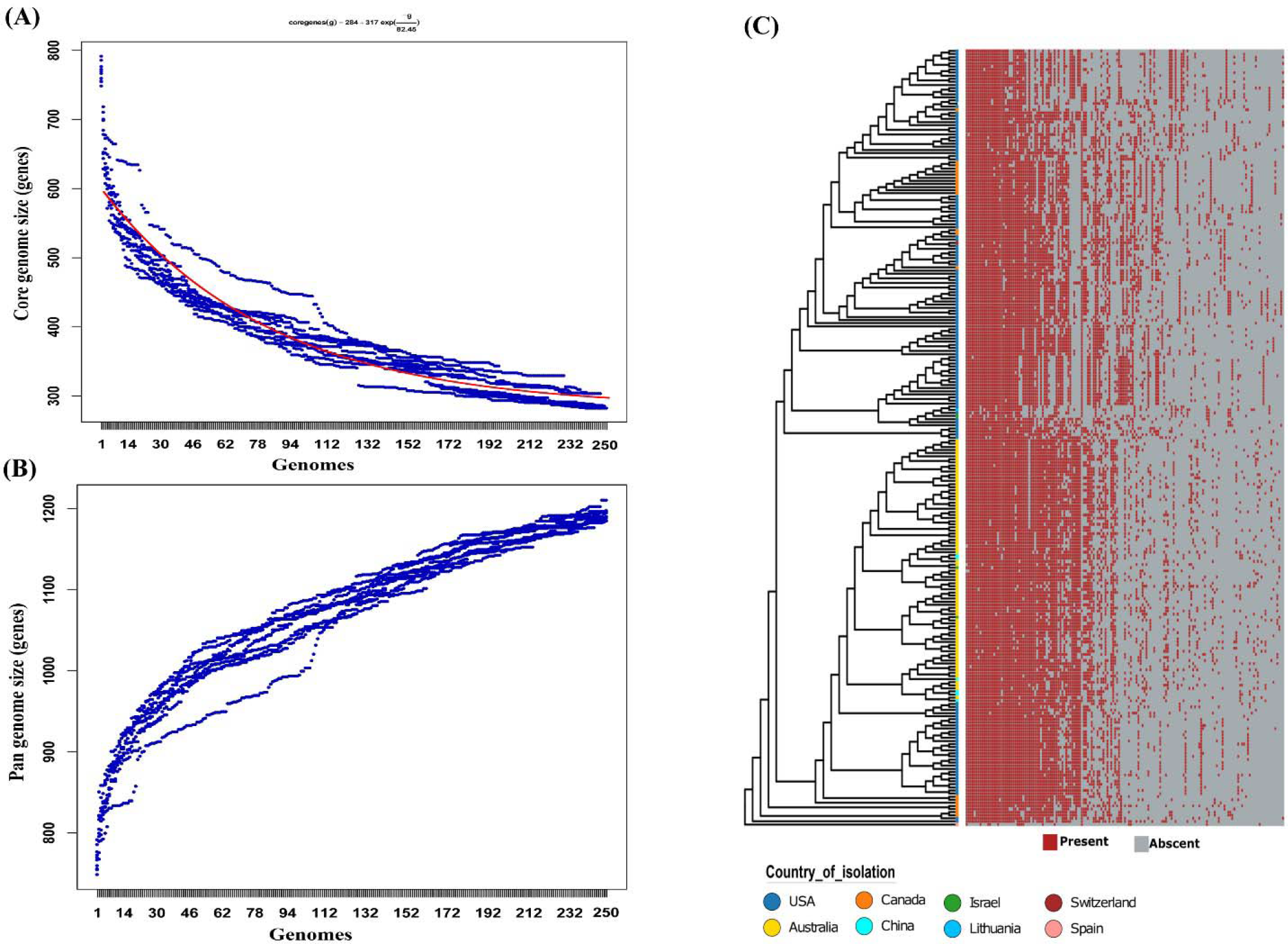
The estimation of core genome and pan genome structure of *M. bovis* isolates. A) & B) represents the core genome (n=283) plot with Tettelin fit and Pan genome (n= 1186) estimated using get_homologues at a query coverage and percentage identity of 75% and 90%, respectively. C) The gene presence-absence matrix of the pangenome from Roary was plotted against the core-genome based phylogeny of *M. bovis*. Genes with a distribution 5% < x < 99% are shown for better clarity.

Apart from this, the ATP-Binding Cassette (ABC) superfamily proteins were also abundant in the core genome. A complete set of genes for an ABC uptake system *i*.*e*., the *oppABCDF* transporter, was conserved across all the *M. bovis* isolates. This system is well known to transport peptides that not only help in cell nutrition [37], but also affects the cell viability [38]. In addition to this, the spermidine/putrescine importer genes *potA, potB* and *potC* were also present in the core genome. Altogether, the dominant subsystems in the core genome were related to protein metabolism, nucleosides and nucleotides, carbohydrates, RNA and DNA metabolism, Genes responsible for fatty acids, lipids and the isoprenoides subsystem were not present, suggesting that *M*.*bovis* depends on the host for their production

### Function-based clustering and its implication for vaccine development

Preventive herd health has become an utmost priority for livestock management and vaccination is a proven backbone for this program. Based on federal regulations, autogenous vaccines must be comprised of strains isolated in conjugation with the animal disease. Therefore, in this study we have included sequence comparisons of ten isolates used as an autogenous vaccine for bison: BB-1, BB-2, BB-3, BB-4, BB-5, BB-6, BB-7, BB-13, BB-14 and BB-17. All were isolated from bison and originated from five different herds in the United States. To assess the degree of functional variation of this blend, we performed function-based clustering (Figure 3). Based on function, we obtained six major clades and interestingly all the Australian isolates clustered together in cluster VI (Figure 3A). All ten vaccine isolates represent four clades; three members each from clades I and II, and two members each from clades III and V (Figure 3A). The clustering pattern of autogenous vaccine isolates suggests that although they are functionally diverse, there are no representatives of certain clades or sub-clades (Figure 3B). In contrast, vaccine isolates within the same lineage sometimes cluster together in close proximity. For example, BB-2/BB-3 in clade V and BB-13/BB-17 in clade III show maximum functional similarity and hence cluster together. Similarly, BB-6/BB-7 (clade I) and BB-5/BB-1 (clade II) are functionally related. Therefore, in order to generate a more broadly effective autogenous vaccine it may be necessary to consider isolates from those clades for which there is no representative strain available in the blend.

**Figure 3:**
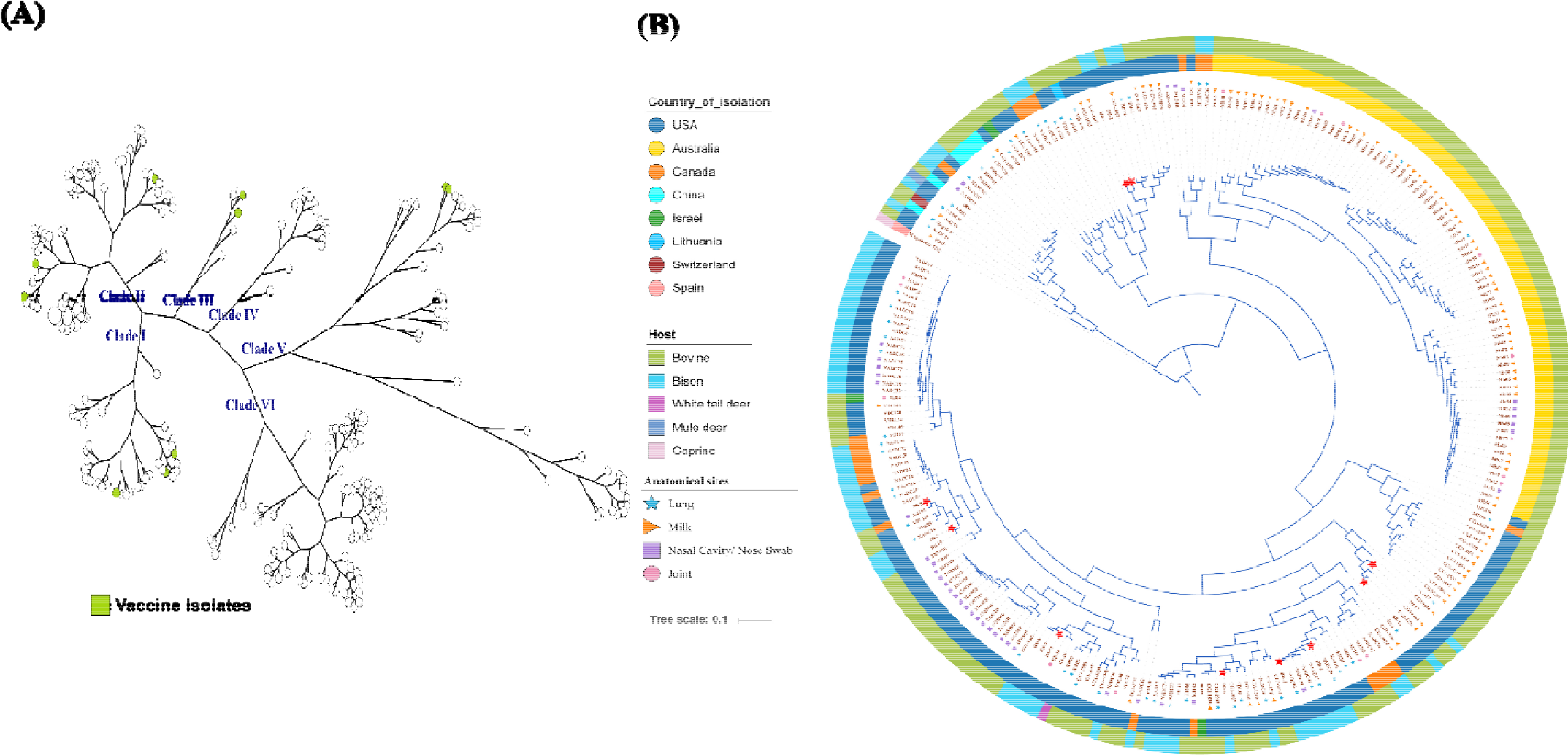
Function-based clustering of *M. bovis* isolates. The kegg matrix was used in MeV to construct the gene tree using Pearson correlation and hierarchical clustering. The tree was visualized using A) GrapeTree (green circles represent the vaccine isolates) and B) iTOL: The outer and inner ring represent the host and country from which they were isolated, respectively. The red star symbols in the branch depicts those strains that are considered for use in autogenous vaccines.

### Vsp dynamics and novel IS elements

A family of *vsp* genes is known to generate a high degree of surface antigenic variation through recombination and thus provides remarkable phenotypic and genetic flexibility [39, 40]. Initially, when we used the 13 *vsp* gene sequences of strain PG45 as a query and searched against the rest of the isolates, none of the isolates were found to harbor the complete set of these *vsp* genes. The most abundant members of this family were *vspJ* (n=73), *vspI* (n=47) and *vspK* (n=37) (Figure 4A). But, when we used the highly conserved 70 bp sequence present immediately upstream of the *vsp* gene start codons as a query, the number of hits was comparatively high. Out of 251 isolates, 29 possessed ≥10 hits, suggesting a high degree of distinctiveness in *vsp* genes at the sequence level (Table 1). In 13 complete genomes, the number of 5’-upstream sequence hits varied from 0 to 14, with 14 in the case of NADC72, and 13 each in NADC81and PG45. No hits were present in the genomes of CQ-W70 and Hubei-1. Among the autogenous vaccine isolates, the number of 5’-upstream sequence hits varied between 2 and 5, suggesting the inclusion of isolates with a high number of *vsp* genes in the autogenous vaccine. Further, the sequence level comparison of these *vsp* genes will be crucial in determining the strain specificity.

**Figure 4:**
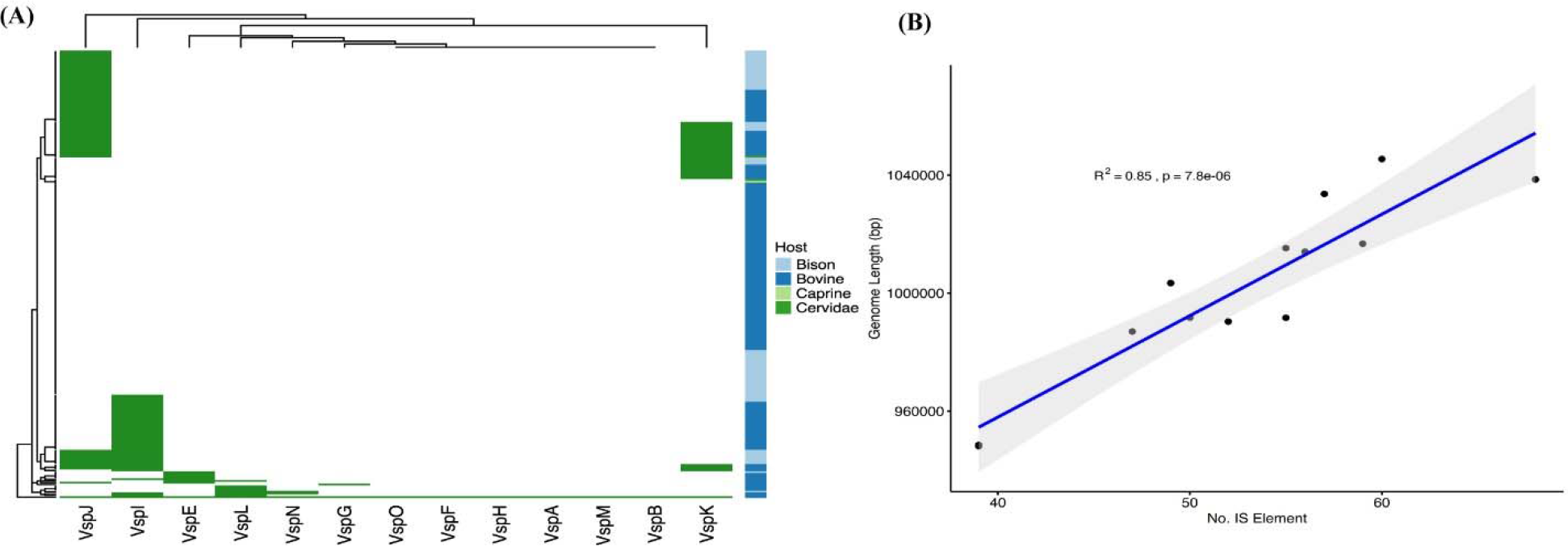
A) The abundance of *vsp* genes across the *M. bovis* isolates used in this study. The *vsp* genes from strain PG45 were used to query the *M. bovis* isolates using a query coverage and percentage identity of 75% and 90%, respectively. B) Scatterplot of genome length and number of IS elements. The p value corresponds to the Pearson Correlation coefficient and the R^2^ is the regression line fit. Shaded area is the 95% confidence interval.

Using a similar approach, we investigated the abundance of ISs in the genomes of the *M. bovis* isolates included in this study. The genome-wide search revealed the presence of a wide variety of IS elements present in the genome. Using match criteria of 70% query coverage and 95% sequence identity, we observed that many isolates have IS elements with a shorter length, *i*.*e*., less than 70% query coverage, but in high abundance. Nevertheless, the average number of IS elements in the complete genomes (N50>900000) and draft genomes (N50>900000) of *M. bovis* isolates is 52.76 and 7.215, respectively. This clearly suggests the sequencing methods and computational tools used are unable to completely resolve the ambiguities of highly repetitive regions [41]. Further, we tested the hypothesis that genome size is correlated with the number of IS elements (N50 ≥ 900000). Our analysis revealed that the number of IS elements found per isolate increases significantly (*p*=7.8e-06) as the genome size increases (Figure 4B). While analyzing the diversity of IS elements, we came across a new 1624-bps element belonging to the IS1634 family of IS elements, which we have named ISMbov9 and the encoded transposase DDE domain (nucleotides 218-1606) is composed of 462 amino acids.

### Gene level prediction of host-associated genes

We applied Scoary to carry out pan-GWAS analysis based on gene presence-absence and the host type, *i*.*e*., bovine versus bison, to predict the genes associated with host types (Table 2). Using this approach, we predicted genes that were significantly associated with host types (*p*<0.05), although the high end of pairwise comparison *p* value range exceeds 0.05 (Table 2). We identified numerous host-associated gene clusters, but most of them are predicted to encode either hypothetical proteins or lipoproteins. Interestingly, a gene coding for variable surface lipoprotein-4 was associated with bovine isolates (p-adjusted value=0.0005) and may play an important role in successful adaptation of *M. bovis* in cattle.

**Table 2:**
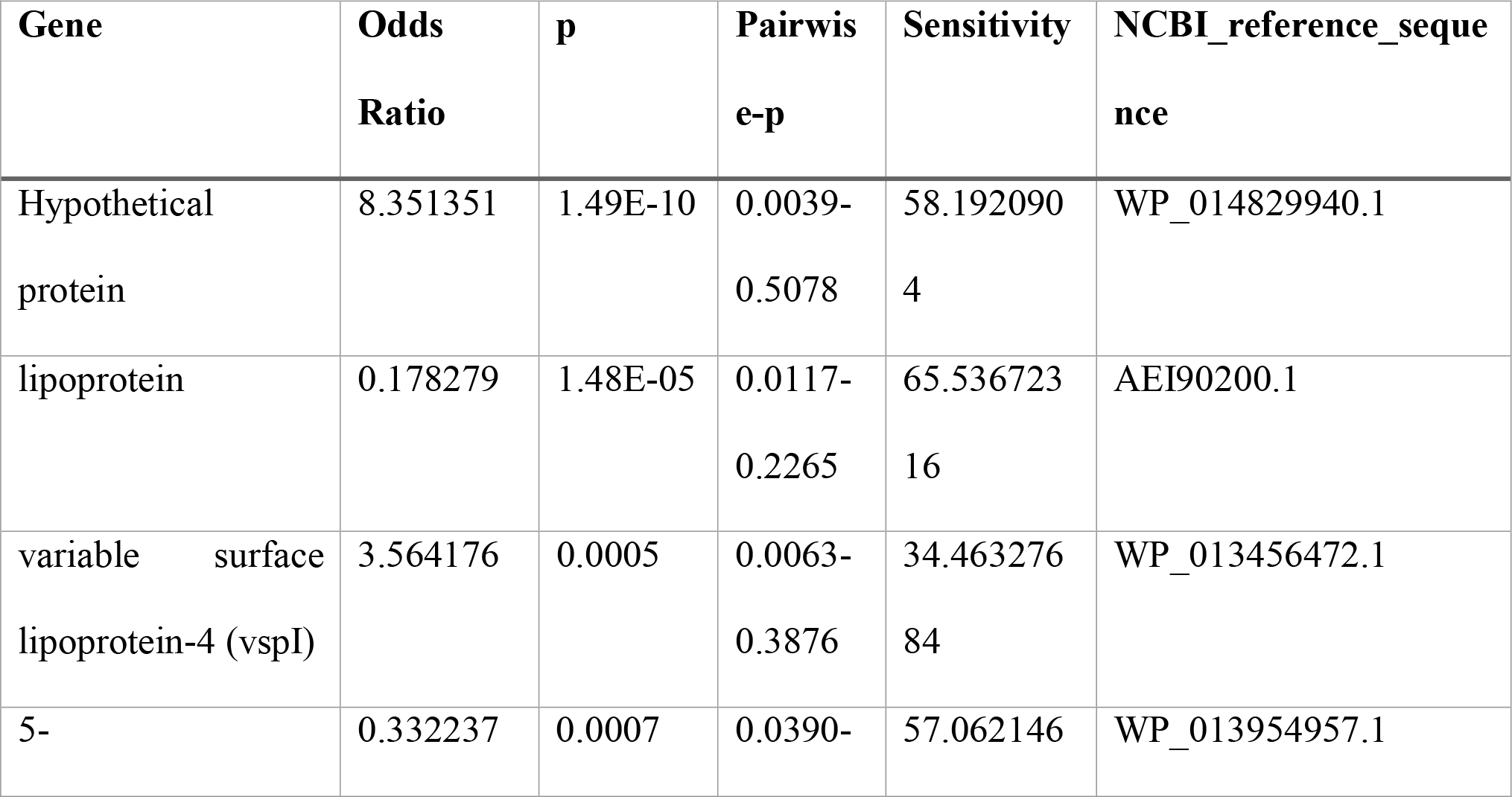
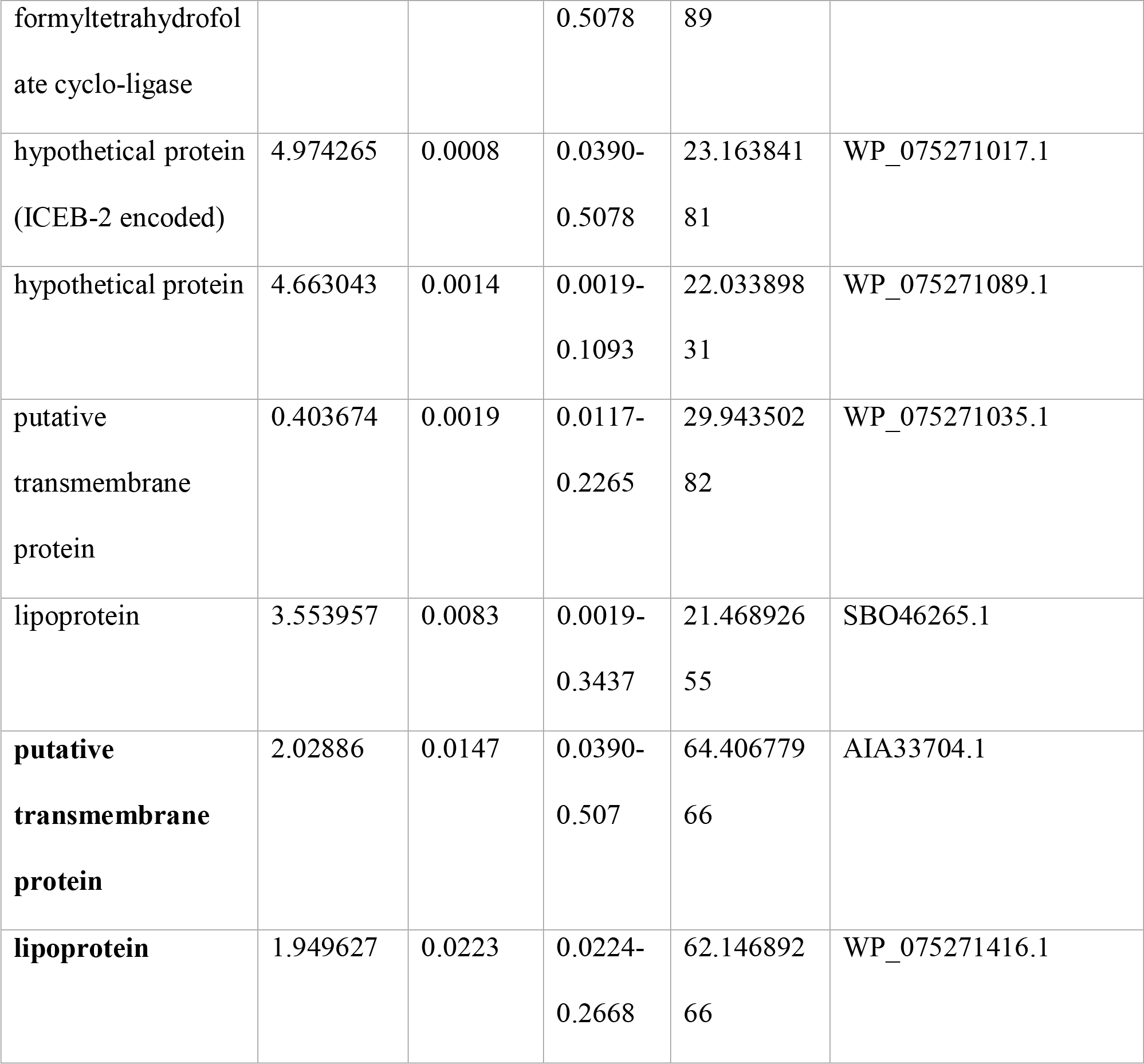
Highest-ranking genes associated with host type (bovine vs bison).

## Discussion

In the present study, we investigated the global population structure of *M. bovis* using the largest genome dataset studied so far. The Australian isolates represent 30.8% of the total population in this study, but their phylogenetic distribution is confined to a specific clade (clade VI) (Figure 1), suggesting a high level of genetic similarity. These results are similar to those of Parker *et al*. [42], who reported that the overall evolutionary changes observed in Australian isolates at the genetic level are minimal despite variations in anatomical sampling sites, geographical location, disease status and time of sample collection. In contrast to this, the American isolates representing 56% of the total population are distributed among five different clades, *i*.*e*., clades I-V; revealing a high level of genomic variation. Similar to this, the Canadian isolates (8.8% of the total population) have a high level of genetic variation and their clustering with American isolates (Figure 1) suggests their origin and spread occurred through the movement of cattle from American herds. The Chinese isolates cluster with Australian isolates, indicating their probable introduction from Australian calves, previously reported using other sequence typing methods [43]. Interestingly, the three Israeli isolates included in this study have two different probable origins; isoaltes 78204 and 88172 appear to derive from Australian calves, while isoalte 87793 is most closely related to isolates from American calves. The overall pattern suggests that the spread of *M. bovis* results from the movement of cattle, similar to the international spread of *M. mycoides* subspecies *mycoides* which causes contagious bovine pleuropneumonia [44, 45].

In spite of the lack of divergence in Australian isolates, the core genome represents only 36.8% of the total genes whereas the number of genes in the pangenome was 1186 as defined here for the genus. The size of the core genome and pangenome fluctuates for any genus depending upon factors such as the percentage identity and query coverage cut-offs values used, the numbers of genomes included, the degree of genomic similarity among the strains and any sampling biases within the taxa [46]. Altogether, the pangenome analysis revealed that the size of the pangenome is increasing steadily with the addition of each genome, suggesting that *M. bovis* has an open pangenome that will expand further by the addition of new genomes.

Our analysis found that the α-enolase protein was conserved in the core genome of *M. bovis*. Although this protein is present in a variety of prokaryotic and eukaryotic organisms [47, 48], it is considered to be a virulence factor in *M. bovis*, playing a role in host cell attachment with the help of host plasminogen [49]. Similarly, two *lplA* genes which have been previously described as potential virulence factors in *M. bovis* [35, 36] are present in the core genome. In *Listeria monocytogenes*, defective LplA protein has been associated with abortive growth along with virulence reduction by 300-folds [50]. Therefore, the presence of genes associated with virulence in the core genome helps the organism in adjusting to the parasitic lifestyle [35, 36, 49, 51]. In addition, the *oppABCDF* transporter and spermidine/putrescine importer genes (*potA, potB* and *potC*) in the core genome help the organism to transport peptides essential for cell nutrition and viability [37, 38], and function in cell proliferation and differentiation, respectively [52, 53].

Our study also describes a functional genomics-based analysis of a multivalent autogenous vaccine comprised of isolates from multiple clades. Using the KEGG annotations pipeline [26], the formulations could be derived by selecting isolates with functional heterogeneity. In the case of *M. bovis*, genotypic differences have been observed among isolates from a single herd and even at different anatomical sites within the same animal [54, 55]. We believe that implementing traditional empirical approaches to screen individual isolates for vaccine development is time-consuming when considering the difficulty of cultivating and maintaining a large number of isolates in the laboratory. Therefore, for a pathogen like *M. bovis*, where the mechanism of pathogenesis is under study, implementing a functional genomics approach could potentially provide new avenues for vaccine development [56].

It has been reported previously that the genetic system for antigenic variation in *M. bovis* is highly complex, mediated through *vsp* genes which undergo dynamic and spontaneous changes in size and expression leading to extensive sequence variations [12, 39, 57-59]. When we analyzed the presence of *vsp* genes in all the isolates using *vsp* genes (n=13) from strain PG45 as a reference, all the isolates evaluated showed the absence of a complete set of *vsp* genes. However, a few genes, namely *vspJ, vspI*, and *vspK*, were detected in 73, 47 and 37 isolates respectively. This suggests that of the *vsp* genes found in type strain PG45, *vspJ* is the most prevalent among other isolates of the species, followed by *vspI* and *vspK*.

In contrast to this, when we used the highly conserved 70 bp sequence present immediately upstream of the *vsp* gene start codons to identify a wider variety of *vsp* genes, we found 29/251 isolatespossess ≥10 sequences, suggesting a high degree of distinctiveness in *vsp* genes at the sequence level. This massive variation in *vsp* sequences defines the vastly extended antigenic potential within the *M. bovis* isolates [57]. The persistence of this pathogen in wide variety of hosts and tissues suggests that it can easily adapt to environmental fluctuations as well as host defense mechanisms with the help of varied antigenic phenotypes [40]. This is in accordance with the results obtained in this study suggesting high degree of distinctiveness at *vsp* genes level. Not only the *vsp* genes diversity, but also the IS elements are playing a crucial role in defining the genome heterogeneity of *M. bovis*. Our results suggest that the IS elements are significantly increasing the genome size of *M. bovis* isolates and their average number in the complete genome was as high as 52.76, suggesting the distribution of these genes across the genomes. During the genome analysis, we identified ISMbov9, a novel insertion element of size 1624 bps. This suggests that the IS elements present in *M. bovis* is still understudied. Therefore, further investigations are needed to analyze the genetic diversity of *vsp* genes in *M. bovis* isolates keeping in mind the possible roles of IS elements. Further, our analysis of genes associated with host type indicates that one *vsp* gene is significantly associated with bovine isolates, suggesting its possible role in adaptation of *M. bovis* in bovine hosts. Altogether, this analysis supports the inclusion of isolates with a high number of *vsp* genes in the autogenous vaccine.

## Conclusions

Despite numerous advances in research pertaining to *M. bovis*, it remains a persistent threat to the cattle and bison industries. *M. bovis* is known to evade the host immune system through extensive antigenic divergence resulting from high recombination efficiency of its *vsp* genes. In addition, the organism is inherently refractory to a wide group of antibiotics due to lack of a cell wall. Therefore, in order to devise successful vaccines it is important to understand the key genomic differences and the extent of diversity among *M. bovis* isolates. In this study, 147 genomes of strains isolated from four different countries were sequenced and combined with publically available *M. bovis* genome datasets in a first-ever large scale comparative study of 251 *M. bovis* genomes. The analysis also focused on the host origin of the isolates to understand the *M. bovis* virulence-host association patterns. Our results revealed high divergence among isolates originating from the USA which clustered into five different clades based on single nucleotide polymorphisms (SNPs). Isolates from Canada, Switzerland, Israel and Lithuania also displayed SNPs similar to US strains suggesting cross-continent transmission of *M. bovis* strains. On the contrary, strains from Australia were found to be minimally divergent and clustered within a single clade with all strains from China and two isolates from Israel. Although sampling bias is evident in the analyzed datasets, with US isolates representing more than half (56%) of the strains, the divergence of US isolates could not be ruled out as one isolate, CG1-1544, clustered with the outgroup *M. agalactiae* 5632 revealing its highest ancestral homology. Our study did not identify any host association patterns of the *M. bovis* strains as isolate from white tail deer (KRB8) clustered with bovine isolates while that from mule deer (KRB1) clustered with the bison isolates NADC98 and NADC97. This indicates the genomic flexibility of *M. bovis* strains for adaptation to different hosts. The analysis of pangenomic trends suggests a large diversity of *M. bovis* that yet remains to be explored. Analysis of genomic heterogeneity among *vsp* genes revealed that out of the 250 isolates, at least 29 harbor more than 10 *vsp* genes. In addition, a large number of IS elements that mediate recombination events was found in all of the *M. bovis* strains and a novel 1624 bp IS element, ISMbov9, was uncovered highlighting an even larger diversity of IS elements within *M. bovis*. Our results revealed that the previously proposed vaccine candidates reported from bison need to be revisited with the now increased *M. bovis* datasets as their potential functional genomic blend failed to cover all the diverged *M. bovis* representatives for development of an effective vaccine.

## Data Availability

Raw genome sequence data for the M. bovis strains used in this study has been submitted under the bioproject PRJNA534329.

## Notes

### Competing Interest Statement

The authors have declared no competing interest.

## References

1. Maunsell FP, Woolums AR, Francoz D, Rosenbusch RF, Step DL, Wilson DJ, Janzen ED: Mycoplasma bovis infections in cattle. J Vet Intern Med 2011, 25:772–783.

2. Razin S, Yogev D, Naot Y: Molecular biology and pathogenicity of mycoplasmas. Microbiol Mol Biol Rev 1998, 62:1094–1156.

3. Rottem S: Interaction of mycoplasmas with host cells. Physiol Rev 2003, 83:417–432.

4. Hale HH, Helmboldt CF, Plastridge WN, Stula EF: Bovine mastitis caused by a Mycoplasma species. Cornell Vet 1962, 52:582–591.

5. Menghwar H, He C, Zhang H, Zhao G, Zhu X, Khan FA, Faisal M, Rasheed MA, Zubair M, Memon AM, et al: Genotype distribution of Chinese Mycoplasma bovis isolates and their evolutionary relationship to strains from other countries. Microb Pathog 2017, 111:108–117.

6. McAuliffe L, Ellis RJ, Miles K, Ayling RD, Nicholas RA: Biofilm formation by mycoplasma species and its role in environmental persistence and survival. Microbiology 2006, 152:913–922.

7. Le Grand D, Solsona M, Rosengarten R, Poumarat F: Adaptive surface antigen variation in Mycoplasma bovis to the host immune response. FEMS Microbiol Lett 1996, 144:267–275.

8. Calcutt MJ, Lysnyansky I, Sachse K, Fox LK, Nicholas RAJ, Ayling RD: Gap analysis of Mycoplasma bovis disease, diagnosis and control: An aid to identify future development requirements. Transbound Emerg Dis 2018, 65 Suppl 1:91–109.

9. Mycoplasma bovis infections [https://www.cabi.org/isc/datasheet/74495#tooverview]

10. Whitaker WR, Shepherd ES, Sonnenburg JL: Tunable Expression Tools Enable Single-Cell Strain Distinction in the Gut Microbiome. Cell 2017, 169:538–546 e512.

11. Guo Y, Zhu H, Wang J, Huang J, Khan FA, Zhang J, Guo A, Chen X: TrmFO, a Fibronectin-Binding Adhesin of Mycoplasma bovis. International journal of molecular sciences 2017, 18:1732.

12. Burki S, Frey J, Pilo P: Virulence, persistence and dissemination of Mycoplasma bovis. Vet Microbiol 2015, 179:15–22.

13. Konstantinidis KT, Tiedje JM: Genomic insights that advance the species definition for prokaryotes. Proc Natl Acad Sci U S A 2005, 102:2567–2572.

14. Wick RR, Judd LM, Gorrie CL, Holt KE: Unicycler: Resolving bacterial genome assemblies from short and long sequencing reads. PLOS Computational Biology 2017, 13:e1005595.

15. Bankevich A, Nurk S, Antipov D, Gurevich AA, Dvorkin M, Kulikov AS, Lesin VM, Nikolenko SI, Pham S, Prjibelski AD, et al: SPAdes: a new genome assembly algorithm and its applications to single-cell sequencing. J Comput Biol 2012, 19:455–477.

16. Wick RR, Judd LM, Gorrie CL, Holt KE: Unicycler: Resolving bacterial genome assemblies from short and long sequencing reads. PLoS Comput Biol 2017, 13:e1005595.

17. Gurevich A, Saveliev V, Vyahhi N, Tesler G: QUAST: quality assessment tool for genome assemblies. Bioinformatics 2013, 29:1072–1075.

18. Seemann T: Prokka: rapid prokaryotic genome annotation. Bioinformatics 2014, 30:2068–2069.

19. Page AJ, Cummins CA, Hunt M, Wong VK, Reuter S, Holden MT, Fookes M, Falush D, Keane JA, Parkhill J: Roary: rapid large-scale prokaryote pan genome analysis. Bioinformatics 2015, 31:3691–3693.

20. Loytynoja A: Phylogeny-aware alignment with PRANK. Methods Mol Biol 2014, 1079:155–170.

21. Page AJ, Taylor B, Delaney AJ, Soares J, Seemann T, Keane JA, Harris SR: SNP-sites: rapid efficient extraction of SNPs from multi-FASTA alignments. Microb Genom 2016, 2:e000056.

22. Stamatakis A: RAxML version 8: a tool for phylogenetic analysis and post–analysis of large phylogenies. Bioinformatics 2014, 30:1312–1313.

23. Li L, Stoeckert CJ, Jr., Roos DS: OrthoMCL: identification of ortholog groups for eukaryotic genomes. Genome Res 2003, 13:2178–2189.

24. Gardner SN, Slezak T, Hall BG: kSNP3.0: SNP detection and phylogenetic analysis of genomes without genome alignment or reference genome. Bioinformatics 2015, 31:2877–2878.

25. Zhou Z, Alikhan NF, Sergeant MJ, Luhmann N, Vaz C, Francisco AP, Carrico JA, Achtman M: GrapeTree: visualization of core genomic relationships among 100,000 bacterial pathogens. Genome Res 2018, 28:1395–1404.

26. Huerta-Cepas J, Szklarczyk D, Heller D, Hernandez-Plaza A, Forslund SK, Cook H, Mende DR, Letunic I, Rattei T, Jensen LJ, et al: eggNOG 5.0: a hierarchical, functionally and phylogenetically annotated orthology resource based on 5090 organisms and 2502 viruses. Nucleic Acids Res 2019, 47:D309–D314.

27. Howe E, Holton K, Nair S, Schlauch D, Sinha R, Quackenbush J: MeV: MultiExperiment Viewer. In Biomedical Informatics for Cancer Research. edited by Ochs MF, Casagrande JT, Davuluri RV. Boston, MA: Springer US; 2010: 267–277

28. Letunic I, Bork P: Interactive tree of life (iTOL) v3: an online tool for the display and annotation of phylogenetic and other trees. Nucleic Acids Res 2016, 44:W242–245.

29. Brynildsrud O, Bohlin J, Scheffer L, Eldholm V: Rapid scoring of genes in microbial pan-genome-wide association studies with Scoary. Genome Biol 2016, 17:238.

30. Altschul SF, Gish W, Miller W, Myers EW, Lipman DJ: Basic local alignment search tool. J Mol Biol 1990, 215:403–410.

31. Fu L, Niu B, Zhu Z, Wu S, Li W: CD-HIT: accelerated for clustering the next-generation sequencing data. Bioinformatics 2012, 28:3150–3152.

32. Rosales RS, Churchward CP, Schnee C, Sachse K, Lysnyansky I, Catania S, Iob L, Ayling RD, Nicholas RA: Global multilocus sequence typing analysis of Mycoplasma bovis isolates reveals two main population clusters. J Clin Microbiol 2015, 53:789–794.

33. Sulyok KM, Kreizinger Z, Fekete L, Janosi S, Schweitzer N, Turcsanyi I, Makrai L, Erdelyi K, Gyuranecz M: Phylogeny of Mycoplasma bovis isolates from Hungary based on multi locus sequence typing and multiple-locus variable-number tandem repeat analysis. BMC Vet Res 2014, 10:108.

34. Register KB, Thole L, Rosenbush RF, Minion FC: Multilocus sequence typing of Mycoplasma bovis reveals host-specific genotypes in cattle versus bison. Vet Microbiol 2015, 175:92–98.

35. Chen S, Hao H, Zhao P, Thiaucourt F, He Y, Gao P, Guo H, Ji W, Wang Z, Lu Z, et al: Genome-Wide Analysis of the First Sequenced Mycoplasma capricolum subsp. capripneumoniae Strain M1601. G3 (Bethesda) 2017, 7:2899–2906.

36. Li Y, Zheng H, Liu Y, Jiang Y, Xin J, Chen W, Song Z: The complete genome sequence of Mycoplasma bovis strain Hubei-1. PLoS One 2011, 6:e20999.

37. Detmers FJ, Lanfermeijer FC, Poolman B: Peptides and ATP binding cassette peptide transporters. Res Microbiol 2001, 152:245–258.

38. Hopfe M, Henrich B: OppA, the substrate-binding subunit of the oligopeptide permease, is the major Ecto-ATPase of Mycoplasma hominis. J Bacteriol 2004, 186:1021–1928.

39. Lysnyansky I, Rosengarten R, Yogev D: Phenotypic switching of variable surface lipoproteins in Mycoplasma bovis involves high-frequency chromosomal rearrangements. Journal of Bacteriology 1996, 178:5395–5401.

40. Lysnyansky I, Sachse K, Rosenbusch R, Levisohn S, Yogev D: The vsp locus of Mycoplasma bovis: gene organization and structural features. J Bacteriol 1999, 181:5734–5741.

41. Treangen TJ, Salzberg SL: Repetitive DNA and next-generation sequencing: computational challenges and solutions. Nat Rev Genet 2011, 13:36–46.

42. Parker AM, Shukla A, House JK, Hazelton MS, Bosward KL, Kokotovic B, Sheehy PA: Genetic characterization of Australian Mycoplasma bovis isolates through whole genome sequencing analysis. Vet Microbiol 2016, 196:118–125.

43. Lysnyansky I, Freed M, Rosales RS, Mikula I, Khateb N, Gerchman I, van Straten M, Levisohn S: An overview of Mycoplasma bovis mastitis in Israel (2004-2014). Vet J 2016, 207:180–183.

44. Fisher J: The origins, spread and disappearance of contagious bovine pleuro-pneumonia in New Zealand. Aust Vet J 2006, 84:439–444.

45. Xin J, Li Y, Nicholas RA, Chen C, Liu Y, Zhang MJ, Dong H: A history of the prevalence and control of contagious bovine pleuropneumonia in China. Vet J 2012, 191:166–170.

46. Livingstone PG, Morphew RM, Whitworth DE: Genome Sequencing and Pan-Genome Analysis of 23 Corallococcus spp. Strains Reveal Unexpected Diversity, With Particular Plasticity of Predatory Gene Sets. Front Microbiol 2018, 9:3187.

47. Sha J, Erova TE, Alyea RA, Wang S, Olano JP, Pancholi V, Chopra AK: Surface-expressed enolase contributes to the pathogenesis of clinical isolate SSU of Aeromonas hydrophila. Journal of bacteriology 2009, 191:3095–3107.

48. Ji H, Wang J, Guo J, Li Y, Lian S, Guo W, Yang H, Kong F, Zhen L, Guo L, Liu Y: Progress in the biological function of alpha-enolase. Anim Nutr 2016, 2:12–17.

49. Song Z, Li Y, Liu Y, Xin J, Zou X, Sun W: α-Enolase, an Adhesion-Related Factor of Mycoplasma bovis. PLOS ONE 2012, 7:e38836.

50. O’Riordan M, Moors MA, Portnoy DA: Listeria intracellular growth and virulence require host-derived lipoic acid. Science 2003, 302:462–464.

51. Pancholi V: Multifunctional α-enolase: its role in diseases. Cellular and Molecular Life Sciences CMLS 2001, 58:902–920.

52. Kashiwagi K, Miyamoto S, Nukui E, Kobayashi H, Igarashi K: Functions of potA and potD proteins in spermidine-preferential uptake system in Escherichia coli. J Biol Chem 1993, 268:19358–19363.

53. Nicolás MF, Barcellos FG, Nehab Hess P, Hungria M: ABC transporters in Mycoplasma hyopneumoniae and Mycoplasma synoviae: insights into evolution and pathogenicity. Genetics and Molecular Biology 2007, 30:202–211.

54. Register KB, Olsen SC, Sacco RE, Ridpath J, Falkenberg S, Briggs R, Kanipe C, Madison R: Relative virulence in bison and cattle of bison-associated genotypes of Mycoplasma bovis. Vet Microbiol 2018, 222:55–63.

55. Bras AL, Suleman M, Woodbury M, Register K, Barkema HW, Perez-Casal J, Windeyer MC: A serologic survey of Mycoplasma spp. in farmed bison (Bison bison) herds in western Canada. J Vet Diagn Invest 2017, 29:513–521.

56. Seib KL, Dougan G, Rappuoli R: The key role of genomics in modern vaccine and drug design for emerging infectious diseases. PLoS Genet 2009, 5:e1000612.

57. Nussbaum S, Lysnyansky I, Sachse K, Levisohn S, Yogev D: Extended repertoire of genes encoding variable surface lipoproteins in Mycoplasma bovis strains. Infect Immun 2002, 70:2220–2225.

58. Behrens A, Poumarat F, Le Grand D, Heller M, Rosengarten R: A newly identified immunodominant membrane protein (pMB67) involved in Mycoplasma bovis surface antigenic variation. Microbiology 1996, 142 (Pt 9):2463–2470.

59. Zou X, Li Y, Wang Y, Zhou Y, Liu Y, Xin J: Molecular cloning and characterization of a surface-localized adhesion protein in Mycoplasma bovis Hubei-1 strain. PLoS One 2013, 8:e69644.

